# Solid state NMR reveals a parallel in register architecture for an infectious recombinant prion

**DOI:** 10.1101/2021.07.20.453078

**Authors:** Manuel Martín-Pastor, Yaiza B. Codeseira, Giovanni Spagnolli, Hasier Eraña, Leticia C. Fernández, Davy Martin, Susana Bravo, Nuria López-Lorenzo, Alba Iglesias, Rafael López-Moreno, Raimon Sabaté, Sonia Veiga, Human Rezaei, Emiliano Biasini, Víctor M. Sánchez-Pedregal, Joaquín Castilla, Jesús R. Requena

## Abstract

Two alternative architectures have been proposed for PrP^Sc^, the most notorious prion: a parallel in register β stack (PIRIBS) and a 4-rung β-solenoid (4RβS). We challenged these two models by measuring intermolecular ^13^C-^13^C dipole-dipole couplings of ^13^CO-labelled Phe residues in a fully infectious sample of recombinant bank vole PrP^Sc^ (recBVPrP^Sc^) using a PITHIRDS-CT solid state NMR (ssNMR) experiment. To our surprise, data strongly support a PIRIBS architecture. However, the mean distance measured (∼6.5 Å) suggests that a minimum of two of the three Phe residues are not perfectly stacked at the canonical ∼5 Å cross-β distance. Additional ssNMR experiments show some local conformational variability of the Phe residues within limits of a relatively high rigidity. The most parsimonious interpretation of our data is that recBVPrP^Sc^ is arranged as a PIRIBS, although additional conformers with alternative architectures cannot be excluded, including a mixture of PIRIBS and 4RβS.

**Author summary:** PrP^Sc^ is the most notorious prion. It is an infectious protein that cuases fatal neurodegenerative diseases in humans and animals. PrP^Sc^ is the aberrant version of a brain protein, PrP^C^. PrP^Sc^ and PrP^C^ have the same prinary structure, but different secondary, tertiaty and quaternary structures. PrP^Sc^ is capable of templating PrP^C^ to convert to the PrP^Sc^ conformation, which is the basis of its capacity to propagate. Two plausible structural models of PrP^Sc^ have been proposed, the four-rung β-solenoid (4RβS) and the parallel in-register β stack (PIRIBS) model. In both cases the driving force of the templating mechanism consists of “sticky” surface β-strands; however, in the PIRIBS model all the β-strands that conform a PrP^Sc^ monomer lie flat on a surface whereas in the 4RβS model they wind in a corkscrew fashion. Here, we analyzed fully infectious recombinant PrP^Sc^ using a solid state NMR technique, PITHIRDS, that allows probing distances between specific labelled amino acid residues. To our surprise (as we have defended the 4RβS model in the past), results clearly show the presence of a PIRIBS structure in our sample.

## Introduction

The scrapie isoform of the prion protein (PrP^Sc^), the most notorious prion, is an infectious protein that propagates by templating its conformation onto units of its alternatively folded conformer, the cellular prion protein (PrP^C^) [1-3]. As it propagates through the brain, PrP^Sc^ causes fatal neurodegenerative maladies such as Creutzfeldt-Jakob disease of humans, scrapie of sheep, bovine spongiform encephalopathy of cattle and chronic wasting disease of cervids [1,2]. Deciphering the structure of PrP^Sc^ is a critical quest in prion research, as the molecular determinants of its propagative capacity and those of the transmission barriers and strain phenomena are encoded in it [2,4]. It has been known for a long time that PrP^Sc^ is an amyloid, and that conversion of PrP^C^ to PrP^Sc^ involves templating of PrP^C^ to unfold/refold on the surface of a PrP^Sc^ amyloid stack featuring unpaired β-strands, “avid” to form fresh -C=O…H—NH-hydrogen bonds with the incoming PrP chain [3,5]. As for the exact architecture of the PrP^Sc^ amyloid, two plausible models have been proposed: a 4-rung β-solenoid (4RβS) [3,5,6] and a parallel in register β stack (PIRIBS) [6,7]. We have supported the 4RβS model, providing experimental evidence militating in its favor, including low-resolution cryo-electron microscopy (cryo-EM) data that led to the construction of a physically plausible 4RβS atomistic model of PrP^Sc^ [3,5,8].

The advent of methods to prepare recombinant PrP^Sc^ [9,10] has opened the possibility of generating the isotopically labelled material required for solid state NMR (ssNMR)-based structural studies of PrP^Sc^. However, the relatively low output of the conversion methods used initially, all based on protein misfolding cyclic amplification (PMCA) techniques [11], has been a limitation, given that relatively large amounts (milligram quantities) of sample are required for these studies. We recently developed a method, protein misfolding shaking amplification (PMSA), which allows facile production of milligram quantities of *bona fide*, infectious recPrP^Sc^ [12]. We used PMSA to generate natural abundance or isotopically labelled bank vole (BV) PrP^Sc^(109I)23-231. This material is highly infectious, with attack rates of 100% and titers of 6.34·10^4^ LD_50_/μg of PrP in TgVole (∼1x) mice [12]. Furthermore, its biochemical and biophysical properties strongly support the notion that its architecture is similar to that of brain-derived PrP^Sc^, with minor differences and structural nuances [12-14], in line with what has been concluded for other recPrP^Sc^ preparations [15,16]. We subjected this material to a ssNMR experiment capable of distinguishing between the two proposed architectures, 4RβS and PIRIBS: PITHIRDS-CT [17], that has been extensively used to probe the basic architecture of amyloids [7,18,19]. The results obtained, surprising to us, are described here, together with additional studies.

## Results

### Further biochemical characterization of recBVPrP^Sc^(109I)23-231

Our PMSA-based procedure to prepare recBVPrP^Sc^(109I)23-231 includes a PK-treatment step to ensure elimination of any non-converted PrP substrate. Such treatment also eliminates the amino-terminal tail ∼23-96 of recBVPrP^Sc^(109I)23-231 and introduces some internal nicks [12]. For simplicity, we will refer to this PK-treated recBVPrP^Sc^(109I)23-231 as recBVPrP^Sc^. We confirmed its pattern of PK-resistant fragments as reported by us previously [12] using SDS-PAGE (Figures 1A, B). Given that bands in the 5-10 kDa region are not well separated in the 4-12% commercial Tris/glycine SDS-PAGE gels that we used previously [12], we subjected the samples to 15% Tris/glycine gels, which allowed a much better separation, visualization and approximate mass assignment of bands in this region. With this system, we detected the previously described fragments with MW of ∼15 and 9.5 kDa, but also a third one of ∼6 kDa which had not resolved from the ∼9.5 kDa band in our previous analyses (Figures 1A, B). Next, we carried out a complete mass spectrometry-based analysis of the samples, which showed an excellent agreement with the SDS-PAGE-based analyses: PK-treated samples were pelleted by centrifugation, and the pellets denatured in 6 M Gdn/HCl and injected into a nanoHPLC coupled to an ESI-TOF detector. Spectra (Figure 1C) showed fragments N97-S231, Q98-S231, and G92-S231 (the ∼15 kDa band seen in the SDS-PAGE gels); fragments N153-S231 and M154-S231 (the ∼9.5 kDa band seen in the gels) and N97-E152, Q98-N153 and Q98-E152 (the ∼6 kDa band in the gels) (Figure 1D). The larger N97-S231 PK-resistant core is reminiscent of Drowsy-type PrP^Sc^ strains [20]. Cleavage at N153 and M154 is also seen in brain-derived PrP^Sc^ albeit at a substantially lower proportion [21,22].

**Figure 1.**
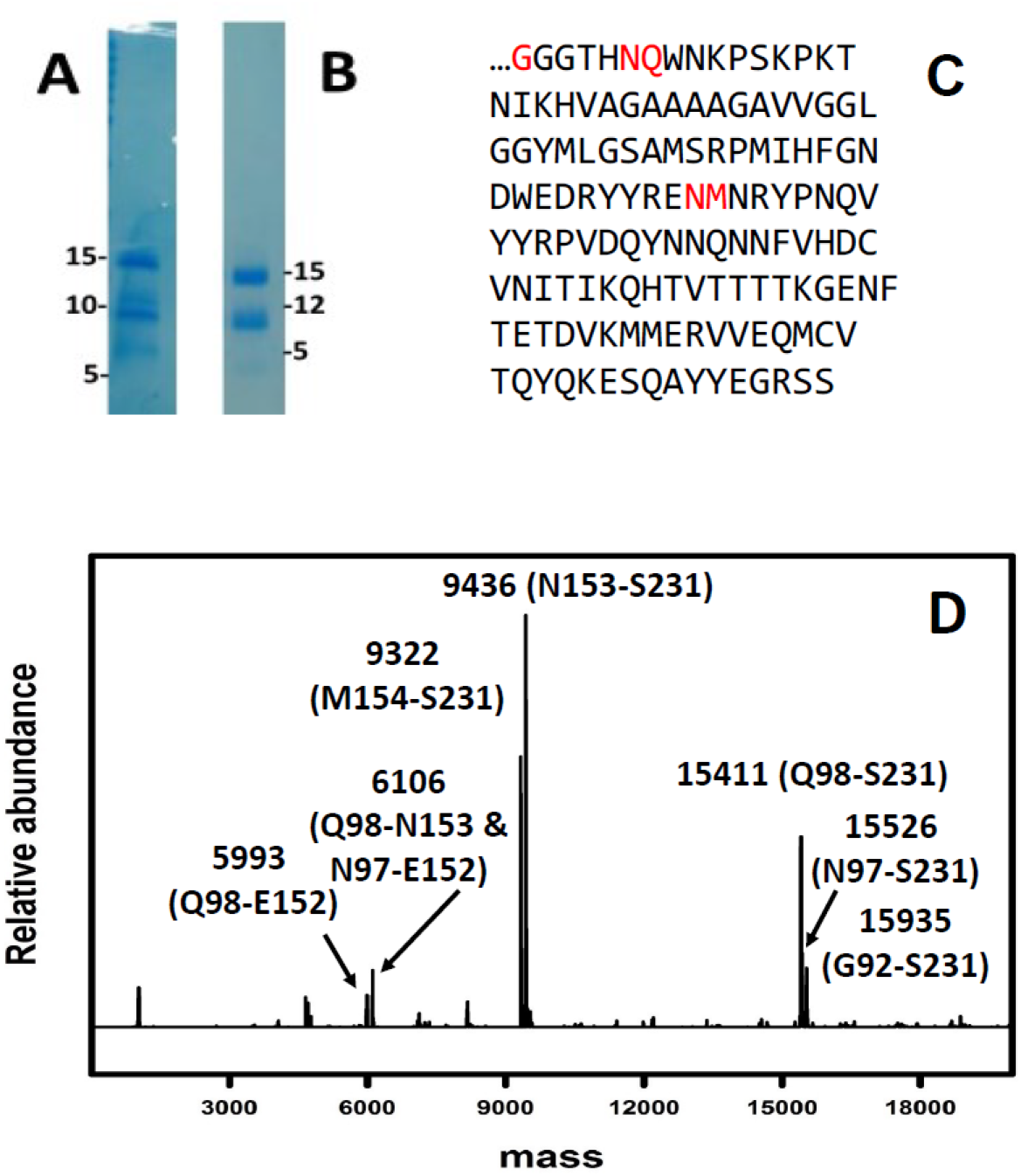
Biochemical characterization of recBVPrP^Sc^. **A)** SDS-PAGE analysis of recBVPrP^Sc^ after treatment with 25 μg/ml PK using commercial 4-12% Tris/glycine gels [12]; given that bands in the 5-10 kDa region are not well separated we subjected the samples to **B)** hand-made 15% Tris/glycine gels, which allowed a much better separation, visualization and approximate mass assignment of bands in this region. Gels were stained with Coomassie blue. **C)** Sequence of recBVPrP^Sc^; PK cleavage sites (*vide infra*) are shown in red. **D)** Mass spectral analysis of the sample revealing three main fragments: N97-S231, Q98-S231, and G92-S231.

### 2D ^1^H-^13^C CP-HSQC spectra of recBVPrP^Sc^ show mostly β-strand-associated signals

We next recorded a 2D ^1^H-^13^C CP-HSQC spectrum of PK-treated uniformly labeled (U-^13^C,^15^N)-recBVPrP^Sc^ (Figure 2). Based on their chemical shifts [23], the majority of signals correspond to residues featuring β-sheet secondary structure (∼85% β-sheet *vs*. ∼15% coil, Figure 2). A proportion of ∼50% β-sheet, ∼50% coil and no α-helix is commonly accepted based on FTIR and CD spectroscopy measurements [3,24,25]. The lower proportion of coil in the NMR spectrum can be a result of the more mobile residues in the flexible loops becoming “invisible” to these NMR experiments, likely a consequence of their dynamics in the μs-ms time scale leading to relative short T_2_ times. We also obtained a variety of ^13^C-^13^C spectra of this sample which will be reported elsewhere. They feature broad signals typical of amyloids [12], also previously seen in similar spectra from PrP non-infectious amyloid samples [18,26].

**Figure 2.**
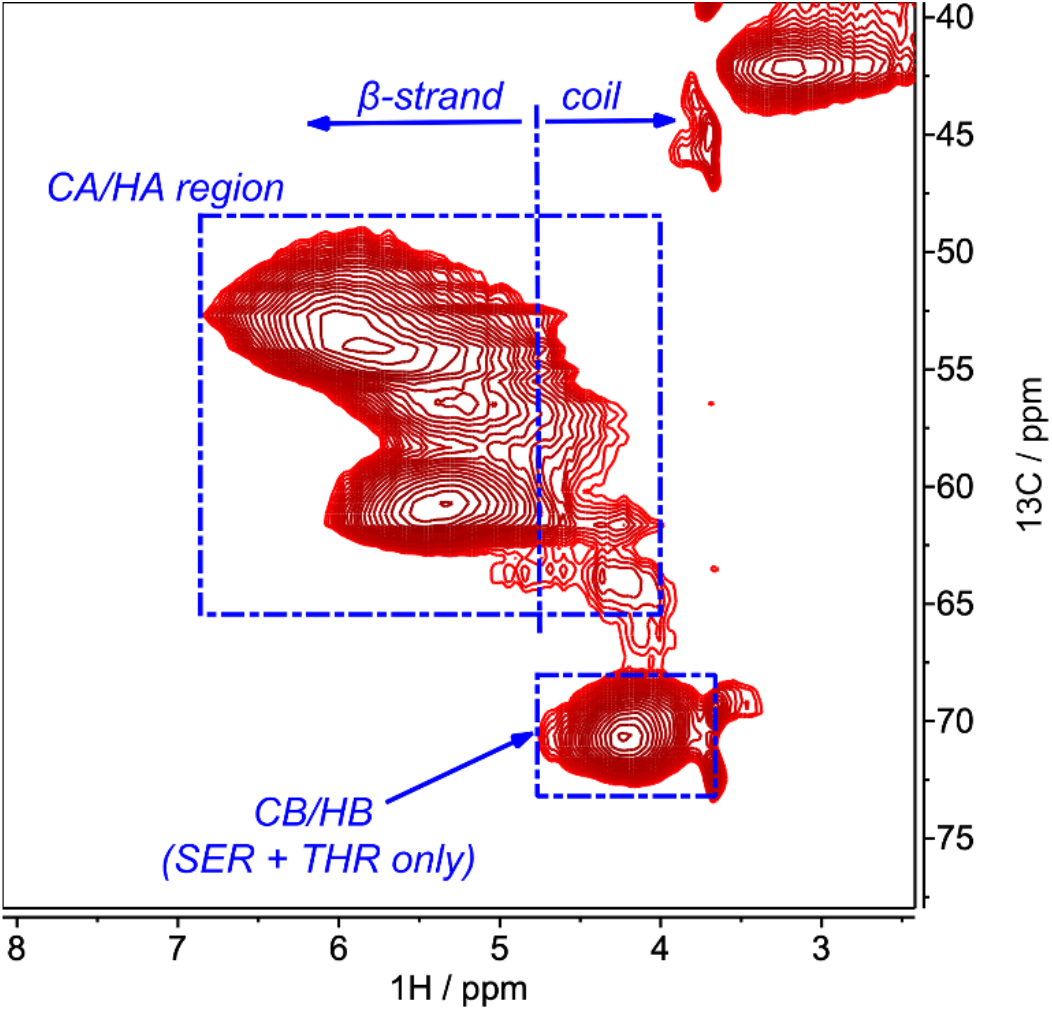
Hα/Cα and Hβ/Cβ regions of the 2D ^1^H-^13^C CP-HSQC spectrum of PK-treated, uniformly labelled (U-^13^C,^15^N)-recBVPrP^Sc^. Choosing a value of 4.6 ppm in the ^1^H axis as the maximum for coil conformation [14] integration of the signals in the resulting areas results in ∼85% β-sheet vs. ∼15% coil, which should be considered cautiously, as the frontier between secondary chemical shifts is fuzzy. However, the majority of signals clearly derive from β-stretches, with just a small fraction arising from connecting loops with intermediate mobility. Signals corresponding to β-strands showed a downfield spread, reaching up to 7 ppm; this, to the best of our knowledge, has not been reported before and might be a consequence of the high packing order and/or effects of the cross-beta CO–HN hydrogen bonds in the chemical shifts of the Hα/Cα resonances.

### PITHIRDS-CT experiments unequivocally show the presence of a PIRIBS recBVPrP^Sc^ conformer

In the absence of a substantial residue-specific assignment, the data obtained from 2D, C-C spectra, while of great descriptive interest, are not sufficient to discriminate between a 4RβS and a PIRIBS architecture. Therefore, we used an experiment capable of distinguishing between both architectures without the requirement of extensive assignment, namely the PITHIRDS-CT developed by Tycko and colleagues [17-19]. During the evolution period of this NMR pulse sequence, rotor-synchronized π pulses that occupy one third of the magic angle spinning (MAS) rotor period are applied on ^13^C to recouple the ^13^C-^13^C dipole-dipole interactions that otherwise average out by MAS. The experiment is repeated for a series of effective recoupling times that modulate the signal intensity. Relaxation and other factors that also modulate signal intensity, in particular during the recoupling period, are kept constant by the use of constant time conditions (CT) for the whole series of experiments [17]. Thus, the curve of ^13^C signal intensity *vs*. effective recoupling time is primarily modulated by the ^13^C-^13^C dipole-dipole interaction and, due to its 1/r^3^ dependence, ^13^C-^13^C distances can be deduced by numerical simulations [7,17-19]. We adapted the pulse sequence and some of the experimental conditions originally developed for a 9.4 T spectrometer to our 17.6 T instrument and 40 kHz MAS. We performed the PITHIRDS-CT experiment on a ^13^CO-Phe labelled recBVPrP^Sc^ sample to probe the intermolecular carbonyl-carbonyl distances. In a canonical PIRIBS architecture, the carbonyl-carbonyl distance between stacked equivalent residues is ∼4.8 Å. In contrast, the intra- and inter-molecular distances between pairs of CO-Phe carbonyl groups in a 4RβS stack are longer than 7 Å [5,19]. We used as controls two amyloids of known architecture: a ^13^CO-Phe labelled recBVPrP non-infectious amyloid sample, known to feature a PIRIBS architecture [18] and a sample of ^13^CO-Tyr labelled HET-s(218-289) fibrils, known to consist of 2-rung β-solenoids [19,27] (Figures S1 and S2). Tycko et al. reported a slightly longer than expected mean distance of 5.0-5.5 Å for the Phe residues in the PrP amyloid [18], explained as a consequence of Phe141 being likely located outside the PIRIBS core. A recent cryo-EM study has confirmed this [28]. In our experiment, the ^13^C carbonyl signal decay confirmed a PIRIBS architecture with a mean Phe-Phe distance of 5-6 Å, in excellent agreement with Tycko’s results (Figures 3 and 4). Also as expected, the signal decay for the HET-s(218-289) prion sample was negligible (Figures 3 and 4), in agreement with the >>7 Å distances between its Tyr residues [19,27]. To our surprise, the ^13^C decay curve of recBVPrP^Sc^ fitted to a mean distance of ∼6-7 Å between Phe residues (Figures 3 and 4). This strongly suggests that the architecture of rec-BVPrP^Sc^ conforms to a PIRIBS, and not to a 4RβS as we had expected [3-5,8].

**Figure 3.**
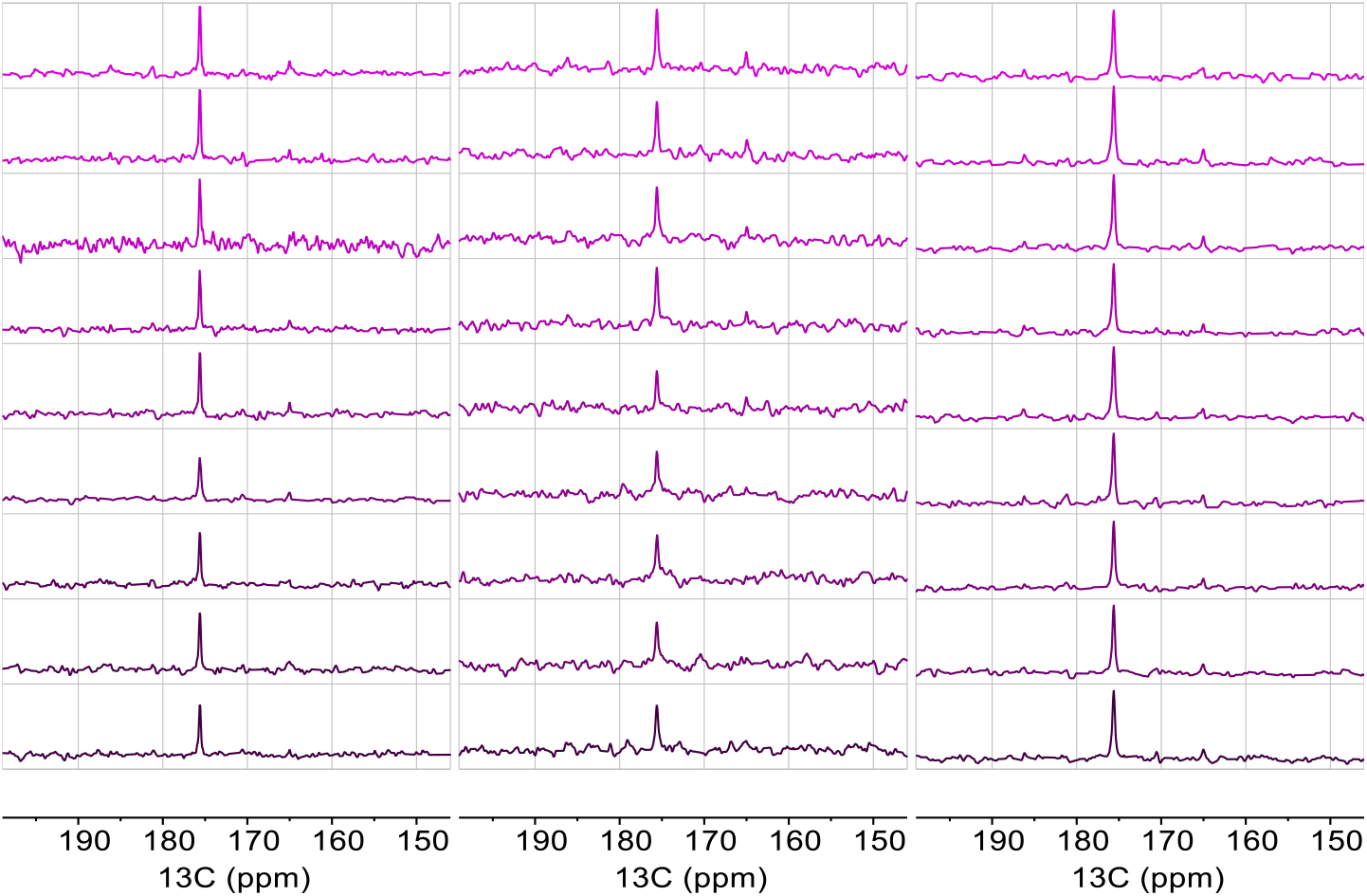
PITHIRDS-CT spectra showing the intensity of the ^13^CO signal. **A)** (^13^CO-Phe)-recBVPrP^Sc^. **B)** (^13^CO-Phe)-recBVPrP(23-231) amyloid. **C)** (^13^CO-Tyr)-HET-s(218-289). From top to bottom, the effective recoupling evolution time is 0.0, 2.4, 4.8, 9.6, 14.4, 19.2, 24.0, 26.4, and 33.6 ms. All spectra of a given sample were acquired with the same number of scans, processed identically and are represented with the same vertical scale. The asterisk denotes a small artefact at the carrier frequency due to pulsed-spin-lock acquisition.

**Figure 4.**
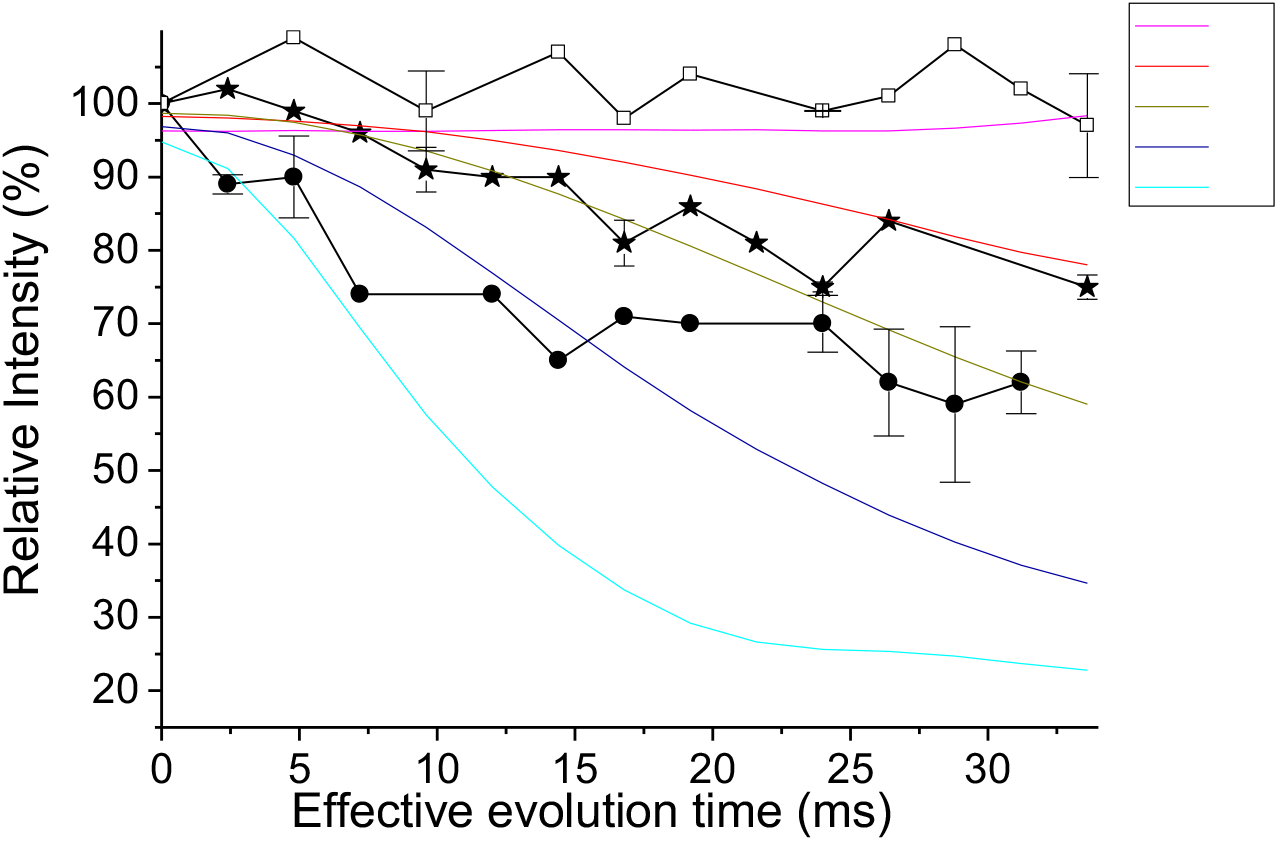
Measurement of intermolecular ^13^C-^13^C dipole-dipole couplings using PITHIRDS-CT. Data are shown after correction of natural abundance ^13^C contributions. Symbols are: • (^13^CO-Phe)-recBVPrP^Sc^, ⋆ (^13^CO-Phe)-recBVPrP(23-231) amyloid, and □ (^13^CO-Tyr)-HET-s(218-289) and represent means of 2 independent experiments (3 for (^13^CO-Phe)-recBVPrP^Sc^), with bars representing standard errors of the mean. Black segments connecting the symbols are drawn to guide the eye. Solid colored lines are simulated curves for specific distances in Å.

### 1D ^13^C CP-MAS and PARIS experiments show overall rigidity in the environments of the three Phe residues of recBVPrP^Sc^

The fact that the measured Phe-Phe distance is longer than ∼5 Å begs for an explanation. We reasoned that at least two of the three Phe residues must be located outside canonical β-strands featuring an ideal PIRIBS architecture, as the measured mean Phe-Phe distance is even larger than that seen in the PrP amyloid, with just one Phe outside the PIRIBS core [18,28]. To comparatively assess the flexibility of our three structures, we measured the peak intensities in 1D ^13^C CP-MAS and PARIS spectra of the samples (Figure 5).

**Figure 5.**
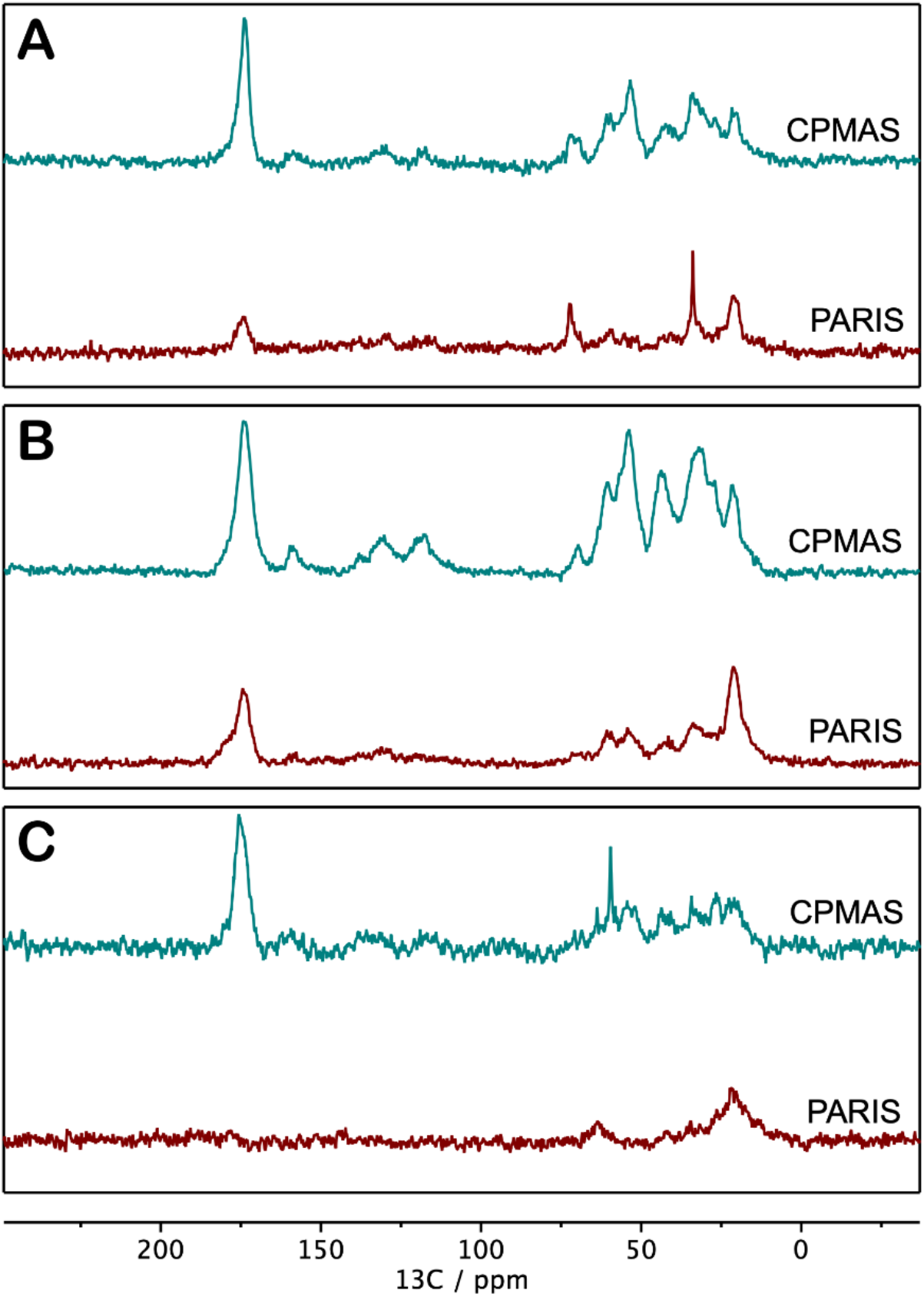
CP-MAS (upper traces) and PARIS (lower traces) spectra of samples: **A)** the (^13^CO-Phe)-recBVPrP^Sc^ prion, **B)** the (^13^CO-Phe)-recBVPrP(23-231) amyloid, and **C)** the fibrillary (^13^CO-Tyr)-HET-s(218-289) prion domain. Aliphatic peaks (0-80 ppm) originate from the ^13^C natural abundance background of the Cα and side chains. The carbonyl band (≈174 ppm) has contributions from the ^13^C labelled residues (**A** and **B**, 3× Phe; **C**, 2× Tyr) and the ^13^C natural abundance background of all the backbone and side chain carbonyl groups. The theoretical contributions of the ^13^C natural abundance to the carbonyl band are: **A:** 38%, **B:** 48%, **C:** 33%.

The PARIS experiment relies on ^1^H-^1^H recoupling and direct polarization from ^1^H to ^13^C. Polarization transfer is effectuated by a hard pulse of just a few microseconds, a duration that is considerably shorter than the usual contact times (*e*.*g*. 0.5 to 5 ms) required for a ^13^C CP experiment [29]. Therefore, PARIS enhances the intensity of nuclei located in moderately flexible stretches characterized by short T_2_ times [30]. Under our experimental conditions, CP provided higher sensitivity with a rigid crystalline sample of glycine with long T_2_ times. In practice, taking as reference the intensities in the CP-MAS spectra, we interpreted that signals that are intense in the PARIS spectrum originate from mobile atoms in flexible stretches, while signals that are very attenuated originate from more rigid stretches (Figure 5).

Resonances of the PARIS spectrum of recBVPrP^Sc^ are heavily attenuated in comparison with the CP-MAS spectrum, reflecting that most of the aliphatic and CO carbons are not experiencing fast dynamics. In particular, the CO resonance drops to <20%. Of the total CO intensity, 62% is due to ^13^CO-Phe spins while the other 38% originates from natural abundance ^13^C background. The CO intensity of the PARIS may originate from those residues (not Phe) displaying some mobility, likely in connecting loops. Similarly fibrillary HET-s, that has a higher percentage of residues in β-sheet stretches, is even more attenuated in the PARIS. In contrast, in the PARIS spectrum of the recBVPrP23-231 amyloid, about 50% of the intensity is conserved both in the aliphatic and carbonyl regions of the spectrum, reflecting residual mobility all over, and agreeing with a higher mobility of Phe141 in this sample [18,28] An extended discussion of these results is provided in the Supporting materials.

### C-C CP-TOCSY spectra show that Phe residues in recBVPrP^Sc^ are in more than one chemical environment

To gather further information on the conformational environment of Phe residues, we prepared recBVPrP^Sc^ with uniformly labelled (U-^13^C,^15^N)-Phe, and recorded its C-C CP-TOCSY spectrum (Figure 6). The cross-peaks in the CA/CB and CB/CA regions are particularly informative: one cross-peak is expected from each one of the Phe residues, unless their chemical shifts completely overlap. Instead of the 3 expected, we detected a minimum of 5 distinct signals in the CA/CB region, which showed better dispersion than the CB/CA region. The simplest explanation is that the side chains of the Phe residues are in two or more different chemical environments (*i*.*e*. conformations). Five distinct signals can be explained if one Phe residue is in three environments (and the other two are in a single environment), or if two Phe are in two environments (and the other one is in a single environment). The fact that these environments are resolved in the C-C TOCSY spectrum indicates that they are either static or in slow exchange in the chemical shift time scale (*i*.*e*., local static disorder). A plausible interpretation is that one or two of the three Phe residues are located in loops featuring alternative conformations for the Phe side chain (as suggested by C-C CP-TOCSY), with very limited dynamic exchange (as suggested by ^13^C-PARIS). These putative alternative conformations do not need to be extremely different: minor packing compactness, hydration or structural accommodation of sidechains may justify the observed changes in the CA and CB chemical shifts. However, a possibility that cannot be ruled out is that two completely different architectures co-exist in separate fibrils (*vide infra*).

**Figure 6.**
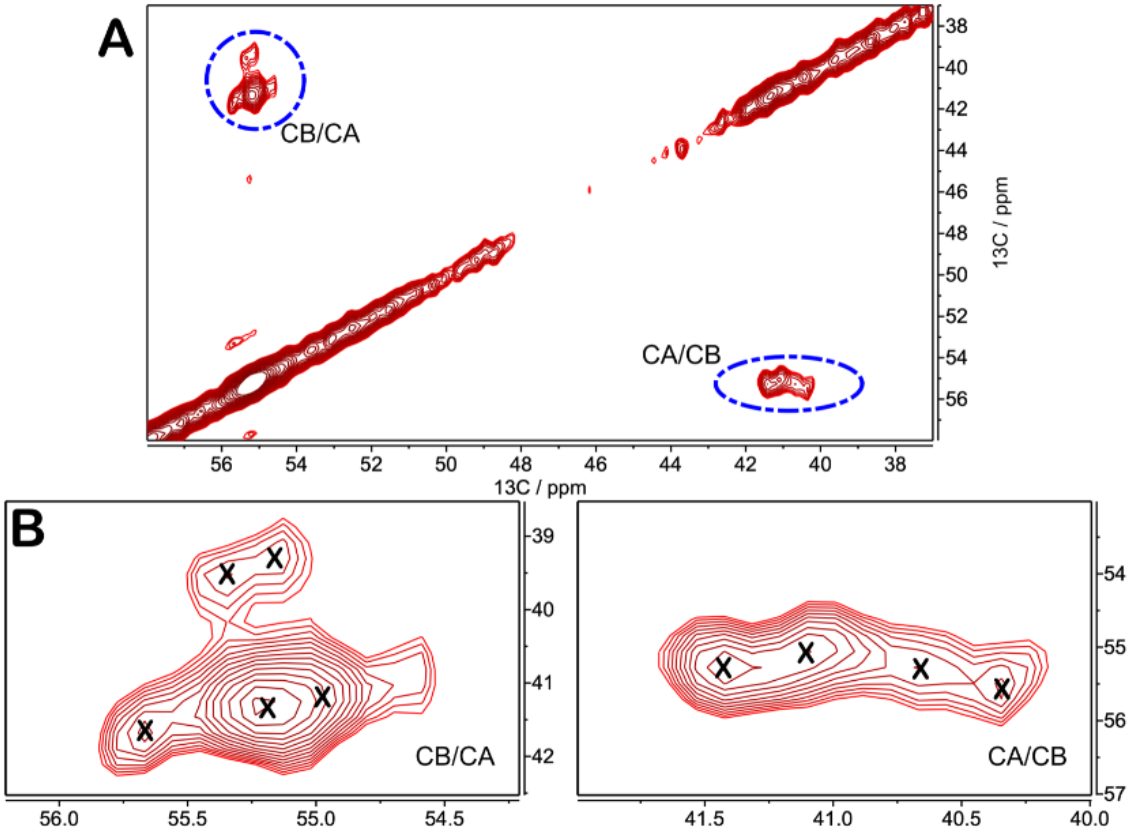
^13^C-^13^C CP-TOCSY with mixing time of 3.55 ms of a (U-^15^N,^13^C-Phe)-recBVPrP^Sc^ sample. **A)** CA/CB region of the spectrum; **B**) and **C)**, expansions.

## Discussion

Our studies unequivocally show that infectious recBVPrP^Sc^ features the characteristics of a PIRIBS, albeit with an unusually long mean distance between stacked Phe residues. One possible interpretation is that two of the three Phe residues in the PK-treated recBVPrP^Sc^ samples are located out of the PIRIBS β-strands, resulting in > 5 Å mean distances. In the non-infectious PrP amyloid, in which Phe141 is out of the PIRIBS core, while Phe175 and Phe198 (BV numbering) are located in β-strands that are part of it [16,28], the mean Phe-Phe distance determined by PITHIRDS is ∼6 Å (Figure 4 and [18]). Therefore, an even higher mean distance of ∼6.5 Å suggests that not one but two of the three Phe residues of recBVPrP^Sc^ are not in a canonical PIRIBS environment. However, as opposed to Phe141 of the non-infectious PrP amyloid, that is located in a relatively flexible albeit not completely disordered environment [18], the putative out-of-β-strand Phe residues of recBVPrP^Sc^ must necessarily be located in relatively rigid environments, as surmised from our PARIS-based analysis (Figure 5). One possibility is their location in short, relatively rigid loops with imperfect cross-β packing and alternative local conformations, connecting *bona fide*, canonical PIRIBS β-strands. Alternatively, the PITHIRDS, CC-TOCSY, CPMAS and PARIS data presented herein could also be explained if our sample comprises a mixture of two components with PIRIBS and 4RβS architectures at a ∼1:2 ratio (Figure S3). It is well known that during the conversion process, structurally diverse PrP^Sc^ assemblies (quasi-species) can be generated, [31] several of which co-exist within a given prion strain [32]. This structural diversity is also responsible for prion adaptation and evolution. However, in our case this would involve two vastly distinct PrP^Sc^ species with exactly the same propagation capacity so as to faithfully maintain their combined biochemical and strain properties over many propagation cycles [12]. This is certainly not very parsimonious, although at this point it cannot be ruled out and further experiments will be required.

While we were finishing these studies, Kraus *et al*. reported on the structure of Syrian hamster 263K PrP^Sc^ resolved by cryo-EM at 3.2 Å resolution [33]. The structure of the main, if not only fibrillary component in the sample is a PIRIBS. Of note, two of the three Phe residues, Phe141 and Phe198, are located in short, packed, apparently rigid loop stretches. Furthermore, Phe198 is near a region that was not resolved due to structural variability [33]. It is tempting to speculate that such structural variability might be the result of alternative conformations of a loop that in the case of recBVPrP^Sc^ might extend to Phe198. While PDB coordinates have not been made available, these features seem compatible with the first interpretation of our data, provided a relatively similar architecture of recBVPrP^Sc^ and brain 263K PrP^Sc^.

All these experimental data run contrary to the notion of a 4RβS PrP^Sc^ conformer. There is however a vast body of experimental evidence supportive of such architecture [3,5,8,34,35], including our own previous data, which will need extensive re-assessment if only PIRIBS PrP^Sc^ exists.

## Materials and Methods

### Production of recombinant protein

RecBVPrP(109I)23-231 was expressed by competent *E. coli* Rosetta (DE3) bacteria harboring pOPINE expression vector containing the wild type I109 bank vole Prnp gene (https://pubmed.ncbi.nlm.nih.gov/29094186). Bacteria from a glycerolate maintained at –80 °C were grown in a 250 ml Erlenmeyer flask containing 50 ml of LB broth overnight at 37 °C and 220 rpm. The culture was then transferred to two 2 L Erlenmeyer flasks containing each 500 ml of minimal medium supplemented with 3 g/L glucose, 1 g/L NH_4_Cl, 1 M MgSO_4_ (1 ml/L), 0.1 M CaCl_2_ (1 ml/L), 10 mg/ml thiamine (1 ml/L) and 10 mg/ml biotin (1 ml/L). For production of uniformly labelled (U-^13^C,^15^N)-PrP, glucose and NH_4_Cl were substituted by (U-^13^C)-glucose and ^15^NH_4_Cl (Cortecnet, Paris) as the sole carbon and nitrogen sources. When the culture reached an OD_600_ of ∼0.9-1.2 AU, isopropyl β-D-1-thiogalactopyranoside (IPTG) was added to induce expression of PrP overnight under the same temperature and agitation conditions. Bacteria were then pelleted, lysed, inclusion bodies collected by centrifugation, and solubilized in 20 mM Tris-HCl, 0.5 M NaCl, 6 M Gdn/HCl, pH = 8. Although the protein does not contain a His-tag, purification of the protein was performed with a histidine affinity column (HisTrap FF crude 5 ml, GE Healthcare Amersham) taking advantage of the natural His present in the octapeptide repeat region of PrP. After elution with buffer containing 20 mM Tris-HCl, 0.5 M NaCl, 500 mM imidazole and 2 M Gdn/HCl, pH = 8, the quality and purity of protein batches was assessed by BlueSafe (NZYTech, Lisbon) staining after electrophoresis in SDS-PAGE gels. Finally, Gdn/HCl was added, to a final concentration of 6 M, for long-term storage of purified protein preparations at –80 °C. For production of (^13^CO-Phe)-PrP and (U-^13^C,^15^N-Phe)-PrP the bacteria were first grown in a 250 ml Erlenmeyer flask containing 50 ml of LB broth overnight at 37 °C and 220 rpm. The culture was then transferred to two 2 L Erlenmeyer flasks containing each 500 ml of minimal medium supplemented with all essential non-labelled L-amino acids except the target amino acid at a concentratiom of 0.1 g/L each, and the labelled target amino acid also at a concentration of 0.1 g/L, and transferred to two 2 L Erlenmeyer flasks. The cultures were incubated at 37 °C, 225 rpm, for 2-3 hours until OD was 0.8 or higher, after which expression was induced by addition of IPTG to a final concentration of 1 mM and the culture was then incubated for 3 hours at 37 °C, 225 rpm. Bacteria were afterwards pelleted and processed as described above.

### Conversion of recBVPrP(109I)23-231 to recBVPrP^Sc^

Conversion was carried out by PMSA as previously described [12]. Briefly, the purified recPrP stored in buffer containing 6 M Gdn/HCl (*vide supra*) was diluted 1:5 in phosphate buffered saline (PBS) and dialyzed against PBS at 1:2000 ratio for 1 h at room temperature. The dialyzed sample was centrifuged at 19000 g for 15 min at 4 °C and the supernatant was used for substrate preparation. The concentration of recPrP in the supernatant was measured spectrophotometrically and adjusted to the working concentration, which was 20 μM, unless otherwise indicated. The protein was then mixed at a 1:9 ratio with conversion buffer (150 mM NaCl, 10 g/L Triton X-100, and 0.5% w/v of dextran sulfate sodium salt from *Leuconostoc spp*. with sizes ranging from 6500 to 10000, Sigma-Aldrich in PBS). The substrate was aliquoted and stored at – 80 °C until required. PMSA was performed by transferring 18 ml of the recPrP substrate to a 50 ml Falcon tube containing 2.8 g of washed 1 mm zirconia/silica beads (11079110Z, BioSpec Products Inc.) and 2 ml of recPrP^Sc^ seed obtained from a previous round of PMSA. The tube was placed in a Thermomixer (Eppendorf) and incubated at 39 °C and 7000 rpm in a continuous mode for 24 h.

### Proteinase K digestion and analysis of digestion products

Samples were digested by addition of PK (Roche) from a concentrated stock solution to a final concentration of 25 μg/ml and incubation at 42 °C for 1 h. The sample was then immediately centrifuged at 19000 g at 4 °C for 30 min, the supernatant was discarded and the pellet resuspended and washed with 1 ml of PBS. After a further 30 min at 19000 g at 4 °C, the supernatant was discarded.

For analysis of PK-induced fragmentation, PK-treated recBVPrP^Sc^ pellets were dissolved in Laemmli buffer and heated at 95 °C for 10 min, followed by SDS-PAGE in home-made 15% Tris/glycine gels or commercial 4-12% Tris/glycine gels (NuPage, Thermo-Fisher). After electrophoresis, gels were stained with BlueSafe Coomassie stain (NZYTech, Lisbon). Alternatively, for mass spectrometry-based analysis, the pellets were resuspended in 50 μl of 6 M Gdn/HCl with 3 pulses (5 s each) of a tip sonicator and incubated at 37 °C for 1 h. TFA was added to a final concentration of 1%. Samples (4 μl) were injected to a micro liquid chromatography system (Eksigent Technologies nanoLC 400, SCIEX) coupled to a high speed Triple TOF 6600 mass spectrometer (SCIEX) with a micro flow source, and equipped with a silica-based reversed phase column Chrom XP C18 150 × 0.30 mm, 3 mm particle size and 120 Å pore size (Eksigent, SCIEX). A YMC-TRIART C18 trap column was connected prior to the separating column, online (3 mm particle size and 120 Å pore size, YMC Technologies, Teknokroma). After sample loading and washing with 0.1% formic acid in water to remove Gdn/HCl and other non-peptide components of the sample, the flow was switched on to the analytical column and separation proceeded at a flow rate of 5 µl/min with a solvent system consisting of 0.1% formic acid in water as mobile phase A, and 0.1% formic acid in acetonitrile as mobile phase B. Peptides were separated over 40 min with a gradient ranging from 2% to 90% of mobile phase B. Data acquisition was performed in a TripleTOF 6600 System (SCIEX, Foster City, CA) using a Data dependent workflow. Source and interface conditions were the following: ionspray voltage floating (ISVF) 5500 V, curtain gas (CUR) 25, collision energy (CE) 10 and ion source gas 1 (GS1) 25. Instrument was operated with Analyst TF 1.7.1 software (SCIEX, USA). Switching criteria was set to ions greater than mass to charge ratio (m/z) 350 and smaller than m/z 1400 with charge state of 2-5, mass tolerance 250 ppm and an abundance threshold of more than 200 counts (cps). Former target ions were excluded for 15 s. The instrument was automatically calibrated every 4 hours using tryptic peptides from PepCalMix as external calibrant. For data analysis, the sample TIC was opened using the Peak View 2.2 software that allows protein reconstruction. The LC-MS Peptide Reconstruct feature uses a peak finding algorithm to identify groups of peaks that form isotope series and charge series. Protein deconvolution was carried out between 800 to 20000 Da.

### Recombinant PrP amyloid fibers

(^13^CO-Phe) and (U-^13^C,^15^N-Phe) recombinant non-infectious BVPrP(109I)23-231 fibers were prepared as described by Torrent *et al*. [36]. Briefly, a 10 ml solution of PrP at 0.6 mg/ml in 50 mM MES buffer, pH 6.0 containing 2.4 M Gdn/HCl was placed in a 15 ml Falcon tube. The tube (arranged horizontally on the plate surface) was incubated with continuous orbital shaking at 30 rpm. (16 mm amplitude) at 37 °C. Fibril formation was monitored using a ThT binding assay: aliquots were withdrawn and diluted into 10 mM sodium acetate buffer, pH 5.0 to a final PrP concentration of 0.3 μM. Then ThT was added to a final concentration of 10 μM and fluorescence measured at λ_ex._ 450 nm, λ_em._ 485 nm. When ThT fluorescence reached the plateau stage, samples were dialyzed in 10 mM sodium acetate, pH 5.0, and collected by ultracentrifugation for 45 min at 228147 g using a Beckman Optima TL100 Ultracentrifuge and TLA-100.3 rotor, and resuspended in 10 mM sodium acetate, pH 5.0. A washing step was performed by repeating the ultracentrifugation and resuspension steps. The final pellet was resuspended in deionized water. For TEM imaging, fibers were adsorbed on Formvar carbon-coated grids, washed with water thrice, negatively stained with freshly filtered 2% uranyl acetate, air-dried and viewed using a JEOL JEM-F200CF-HR electron microscope.

### HET-s fibers

The His_6_-HET-s prion forming domain Het (218-289) [37,38] was expressed in BL21 *E. coli* and purified using NTA affinity chromatography as described for PrP. The protein buffer was exchanged to 200 mM acetic acid (pH = 2.5) using a PD-10 gel filtration column. Protein concentration was adjusted to 200 μM and an equal volume of 1 M Tris/HCl, pH 8 was added. The resulting sample was incubated at 37 °C with shaking at 1400 rpm in a Thermomixer for 24 h. A visible bulky sediment was visible at the end of the incubation period, which was collected by centrifugation and resuspended in the desired volume of deionized water. The sample was imaged by TEM as described for the PrP amyloid.

### Solid-State NMR measurements (ssNMR)

#### Sample preparation

(U-^13^C,^15^N)-recBVPrP^Sc^ solution, as obtained by PMSA, was treated with 25 μg/ml of PK (*vide supra*). The sample was then centrifuged at 9000 g for 1 h at 4 °C in 85 ml OAK polycarbonate (Nalgene) tubes using a FiberLite F15-6×100 rotor (Piramoon Technologies, Inc.) in a Beckman Sorvall Legend XTR/230V ultracentrifuge. The supernatant was carefully removed, and the pellet washed with milliQ water, centrifuged under the same conditions for additional 15 min and supernatant removed to obtain the final pellet. The sample was then loaded into a 1.3 mm rotor for ssNMR measurement using the Bruker solid sample preparation kit. The final pellet was resuspended in 50 μl water by repeated pipetting and transferred to the loading funnel with the 1.3 mm rotor inserted. This assembly was placed in a home-built desiccation chamber containing CaCl_2_ until all the water evaporated, leaving thin PrP^Sc^ scales around the border of the 1.3 mm rotor. Using the Bruker loading rod, the scales were carefully introduced and compacted into the rotor, together with 2 μl of milliQ water to obtain a hydrated sample. The rotor was capped and placed in the solid NMR probe. Solid-state NMR experiments were measured at 278 K in a Bruker NEO 17.6 T spectrometer (proton resonance 750 MHz and ^13^C resonance 188 MHz) equipped with a ^1^H/^13^C/^15^N triple resonance solid probe for 1.3 mm zirconia rotors and an available range of MAS rates from 8 to 67 kHz.

The spectrometer control software was TopSpin 4.0. All spectra were processed with MestreNova v14.0 (Mestrelab Research Inc.). Carbon-13 chemical shifts were referenced to the CA signal of solid glycine at 43.5 ppm. Nitrogen-15 chemical shifts were referenced to the ^15^N peak of a solid ^15^N labelled sample of glycine at 35.0 ppm. Proton chemical shifts were referenced to the intense water peak at 4.7 ppm.

1D ^13^C cross-polarization spectra (^13^C CPMAS) were measured at a MAS rate of 40 kHz (pulse program *hC*.*cp* of the Bruker library) with an inter-scan delay (d_1_) of 2 s. Cross-polarization was applied during 3 ms with a constant carbon field strength of 66.6 kHz; the power on the ^1^H nucleus was linearly ramped from 70% to 100% with a peak field strength of 152 kHz. Heteronuclear decoupling during FID acquisition was performed with SPINAL-64 with a proton field strength of 170 kHz. The spectrum was acquired with 1500 scans.

1D ^13^C PARISxy^[25]^ direct polarization spectra (^13^C PARIS) were measured at a MAS rate of 40 kHz. At the end of the inter-scan relaxation delay, immediately before the ^13^C excitation pulse, a train of PARISxy 12.5 μs pulses with 24 kHz field strength were applied to introduce modulation sidebands at half the rotation rate, causing ^13^C enhancement. The pre-scan delay (d_1_) was set to 0.5 s, which is followed by PARISxy irradiation during 3 s. The ^13^C excitation pulse had a tilt angle of 90° and was applied with a B_1_ field strength of 62.5 kHz. Heteronuclear decoupling during acquisition of the FID was performed with SPINAL-64 with a proton field strength of 170 kHz. The spectrum was acquired with 1500 scans.

2D ^1^H-^13^C CP-HSQC spectra were measured at 50 kHz MAS (pulse sequence *hCH2D*.*dcp* of the Bruker library). The ^1^H and ^13^C carriers were placed at 2.27 and 80 ppm, respectively. The spectral widths in the ^1^H and ^13^C dimensions were 19.6 and 175 ppm, respectively. The initial proton/carbon cross polarization contact time was 0.7 ms and the proton contact pulse power used an ascending linear ramp of 50%. The final carbon/proton cross polarization contact time was 0.4 ms and the proton contact pulse power used a descending linear ramp of 50%. MISSISSIPPI water suppression pulses were applied on proton at 15 kHz during 7.66 ms. The number of complex points acquired in the t_2_ and t_1_ dimensions were 588 and 300, respectively. The FID was acquired under heteronuclear decoupling at 20 kHz with WALTZ-16 for both ^13^C and ^15^N nuclei. The inter-scan delay was 1.4 s and the number of scans per t_1_ value was 16.

PITHIRDS-CT spectra were acquired on ^13^CO selectively labelled samples at MAS 40 kHz with sensitivity enhancement using Pulse-Spin-Locking (PSL) acquisition [39]. The conditions for the initial cross-polarization step were identical to the ^13^C-CPMAS spectrum described above. The Constant-Time (CT) evolution period with ^13^C PITHIRDS dipolar recoupling was applied for a total of 33.6 ms, as defined by Tycko [19]. It comprises cycles of sixteen rotor periods occupied with 180° pulses of 8.33 μs duration (τ_r_ /3) with displacements in time of 0, τ_r_/3 and 2τ_r_/3, where τ_r_ is the rotor period duration. A series of effective dipolar evolution periods were explored in different spectra ranging from 0 to the maximum of 33.6 ms. During the PITHIRDS-CT sequence and acquisition periods, ^1^H TPPM decoupling at 110 kHz field strenght was applied. Carbon-13 PSL π pulses had a duration of 7 μs and were applied with 20 rotor cycles per echo. The relaxation delay (d_1_) was set to 4 s and each spectrum was acquired with 4096 scans in 4 h 36 min. Data analysis included natural abundance correction for the ^13^C background. PITHIRDS-CT curves were simulated with in-house software written in Fortran, gently provided by Robert Tycko (Laboratory of Chemical Physics, NIDDK, NIH, Bethesda, USA). The simulation of PITHIRDS-CT intensity curves was performed for a system of five ^13^C atoms arranged linearly at distances of 4, 5, 6 and 7 Å.

2D ^13^C-^13^C-TOCSY [40] was acquired for sample (U-^13^C,^15^N-Phe)-recBVPrP^Sc^ at 16.9 kHz MAS. The initial cross-polarization was applied during 2 ms with a constant carbon field strength of 118 kHz; the power on the ^1^H nucleus was linearly ramped from 50% to 100% with a peak field strength of 202 kHz. After t_1_ evolution, spin-lock was applied during 3.55 ms with continous wave pulses simultaneous on ^13^C and ^1^H with field strengths of 51.4 and 100 kHz, respectively. In both ^13^C dimensions the center was placed at 100 ppm and the spectral width was 221 ppm. Heteronuclear decoupling during FID acquisition was performed with SPINAL-64 with a proton field strength of 110 kHz. The indirect dimension was acquired with 300 complex points in t_1_ with the States-TPPI acquisition mode and 833 points in t_2_. The number of scans per t_1_ increment was 96 and the relaxation delay (d_1_) was 2 s. The total measurement time was 16 h 25 min.

## Acknowledgements

We thank Robert Tycko, Laboratory of Chemical Physics, NIDDKD, NIH, for advice and sharing with us PITHIRDS sequence scripts, and Patricia Piñeiro (CIC-BIOGUNE) and María José Pazos (Electron and Confocal Microscopy Core facility, USC) for excellent technical assistance.

## Supporting information

**Figure S1.**
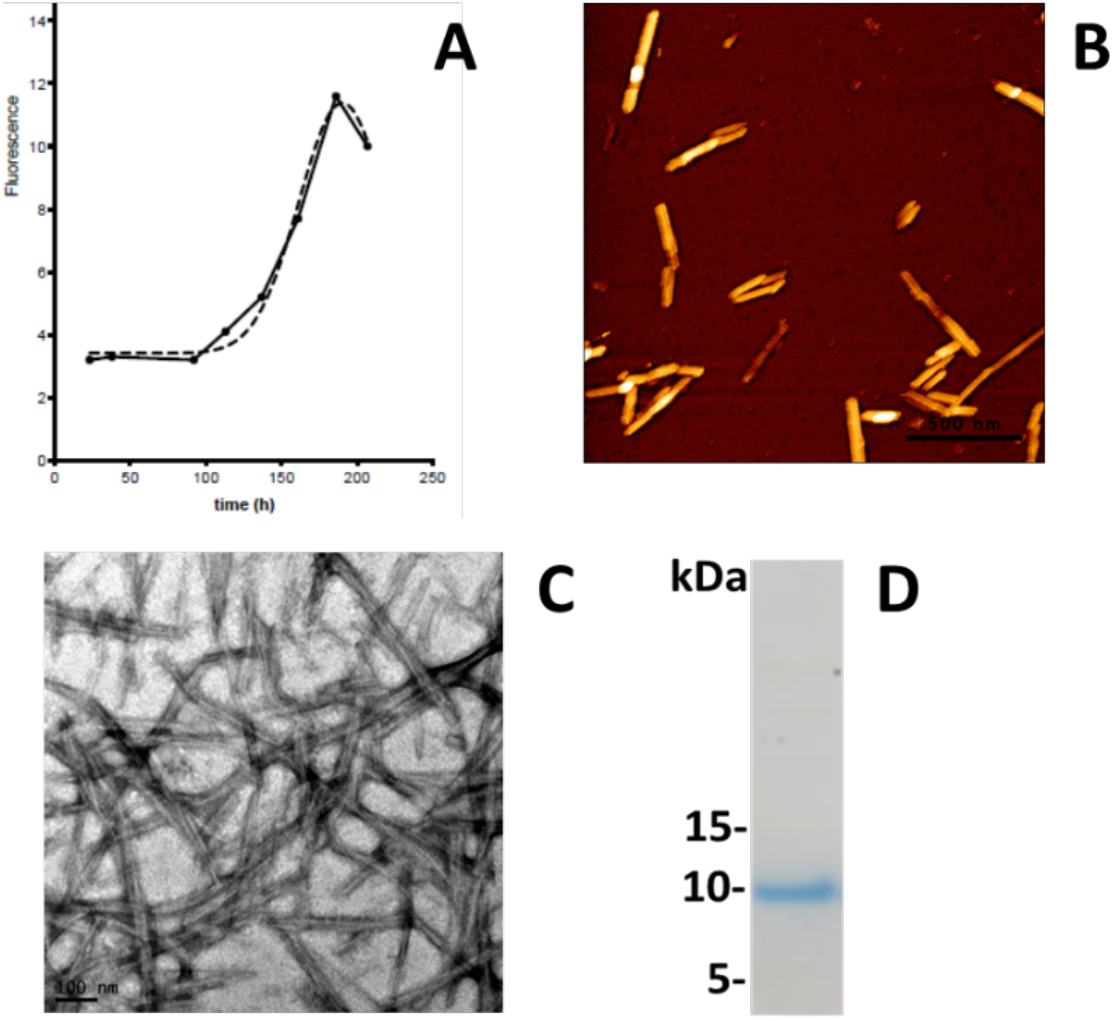
Characterization of the non-infectious recBVPrP(109I)23-231 amyloid sample. **A)** Thioflavin T (ThT) fluorescence. **B)** Atomic force microscopy. **C)** Negative stain Transmission Electron Microscopy (TEM). **D)** Partial PK resistance.

**Figure S2.**
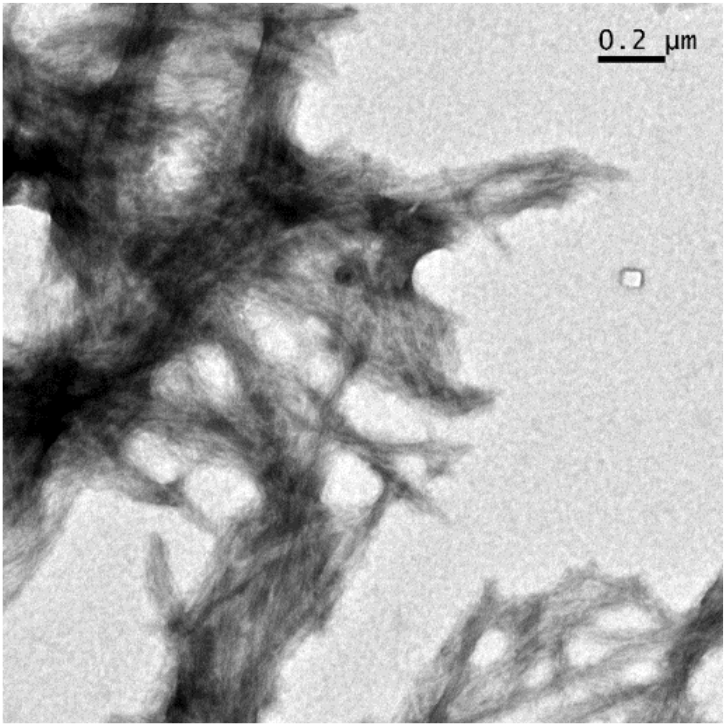
Negative stain TEM image of the fibrillar HET-s(218-289) prion domain sample.

**Figure S3.**
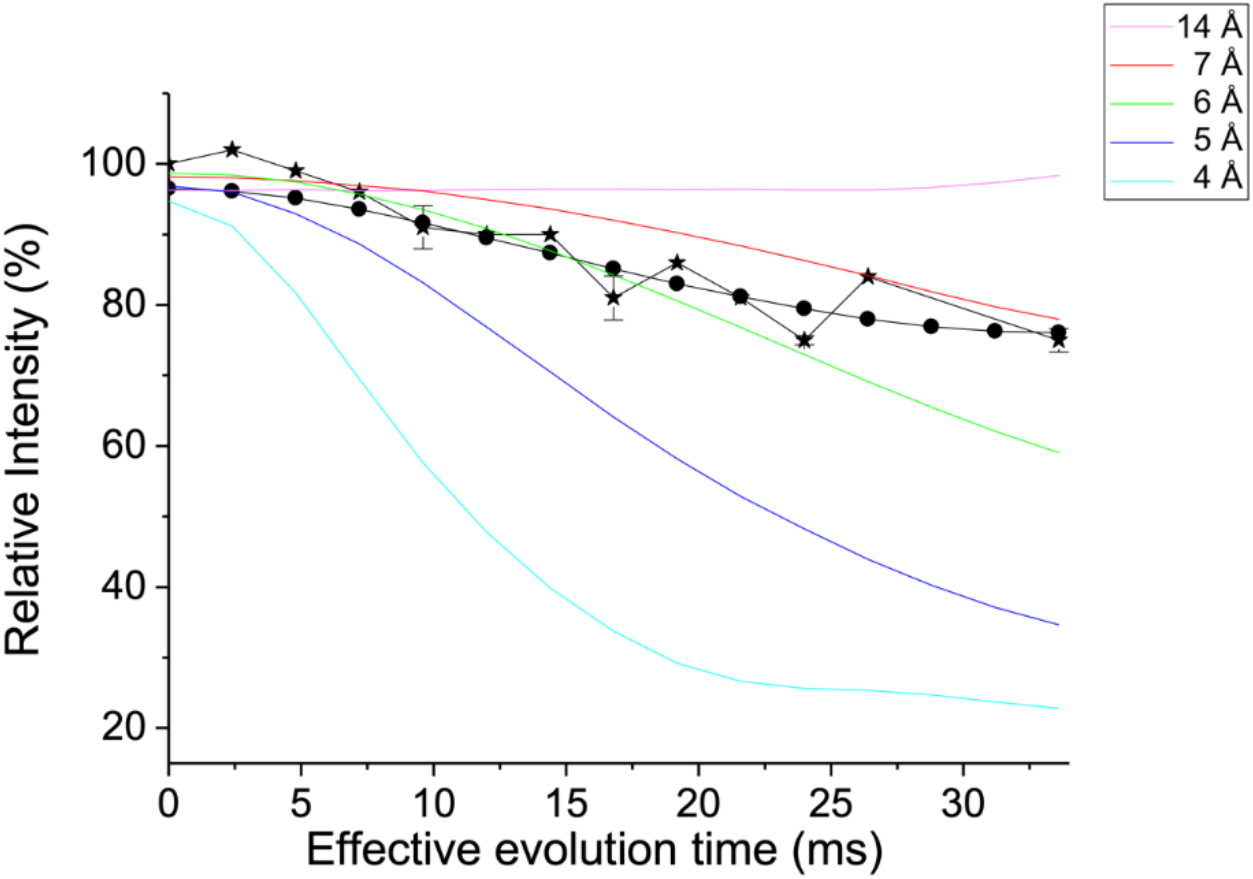
Measurement of intermolecular ^13^C-^13^C dipole-dipole couplings using PITHIRDS-CT. Symbols are: ⋆ *experimental* (^13^CO-Phe)-recBVPrP^Sc^ PITHIRDS-CT intensities with correction of the natural abundance, and • *simulated* averaged PITHIRDS-CT intensities for a model of two structures with no exchange among them consisting in a mixture of 65% of 4RβS and 35% PIRIBS. Black segments connecting the symbols are drawn to guide the eye. Solid colored lines are simulated curves for a linear arrangement of 5 atoms separated by the given distances (4, 5, 6, 7 and 14 Å), as indicated in the main text. Experimental values (⋆) represent the mean of two independent experiments. Simulated points (•) were calculated as the weighted average of two PITHIRDS-CT simulations carried out for a linear arrangement of 5 atoms separated by distances of 14 Å and 5 Å. The weighting factors were manually adjusted for fitting the experimental curve in steps of 5% and the best fit was obtained with factors 0.65 and 0.35, respectively. The intensities simulated for the distance of 5 Å are consistent with the canonical 5 Å ^13^CO-^13^CO cross-β distance characteristic of a PIRIBS structure. The intensities simulated for the distance of 14 Å were very close to those experimentally obtained in Figure 3 (main text) for (^13^CO-Tyr)-HET-s(218-289), which is known to adopt a 2RβS structure.

### Supporting Discussion: Comparison of 1D ^13^C CP-MAS and PARIS spectra

The carbonyl peak (δ_C_ ≈ 174 ppm) of the CP-MAS spectrum of (^13^CO-Phe)-recBVPrP^Sc^ originates from the ^13^C labeled atoms of the three Phe residues plus a contribution from the ^13^C natural abundance background of all the backbone and side chain carbonyl groups. The intensity of the latter contribution can be estimated with the *natural abundance* factor *f* (equation S1, Table S1). All peaks outside of the carbonylic region (δ_C_ ≈ 0-167 ppm) originate from the (unlabeled) ^13^C natural abundance background of the backbone Cα and side chain carbons.

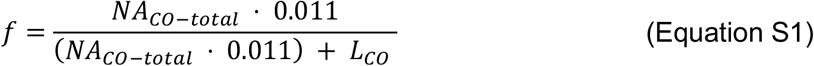

**Table S1.** Contribution (*f*) of the ^13^C natural abundance background to the carbonyl signal of each protein sample analyzed (*L*: Labelled. *NA*: Natural Abundance).

| sample | $L_{CO}$ | $NA_{CO}$-<br>backbone | $NA_{CO}$-<br>sidechain | $NA_{CO}$-<br>total | $f$ |
| --- | --- | --- | --- | --- | --- |
| **A** ($^{13}\text{CO}$-Phe)-recBVPrP<sup>Sc</sup> | 3 | 135 | 35 | 170 | 0.38 |
| **B** ($^{13}\text{CO}$-Phe)-recBVPrP(23-231) amyloid | 3 | 208 | 44 | 252 | 0.48 |
| **C** ($^{13}\text{CO}$-Tyr)-HET-s(218-289) | 2 | 73 | 16 | 89 | 0.33 |

$L_{CO}$ : number of residues with $^{13}\text{C}$ labeled CO.
$NA_{CO}$ -backbone: number of backbone CO at $^{13}\text{C}$ natural abundance.
$NA_{CO}$ -sidechain: number of side chain CO at $^{13}\text{C}$ natural abundance (residues Q, E, N and D).
$NA_{CO}$ -total : $NA_{CO}$ -backbone + $NA_{CO}$ -sidechain

Our goal with the PARIS experiments was to determine if the labeled CO atoms of prion recBVPrP^Sc^ (sample **A**, (^13^CO-Phe)-recBVPrP^Sc^) undergo some relatively fast motion that may explain the signal decay observed in the PITHIRDS-CT experiment. As consequence of our choice of experimental NMR parameters, compared to CPMAS, signals in the PARIS spectrum are enhanced for mobile atoms, while signals from rigid stretches disappear or are very attenuated. The CO signal of sample **A** is 4-fold weaker in the PARIS spectrum than in the CP-MAS spectrum (Figure 3-A), which indicates that, globally, the CO atoms are not very mobile. It is unclear whether the CO signal in PARIS arises from the selectively labeled residues or from the natural abundance background.

In order to support this qualitative interpretation, we compared the signal attenuation of the CO with the other signals of the same PARIS spectrum. Clearly, the attenuation of the CO is more pronounced than the attenuation of the sidechains (some caution should be exercised as the number of protons attached or in close proximity to the majority of carbons of the sidechains is larger than for the CO, *i*.*e*. not only motion counts). We did the same kind of comparison with two control samples: the (^13^CO-Phe)-recBVPrP(23-231) amyloid (sample **B**) and the (^13^CO-Tyr)-HET-s(218-289) prion domain (sample **C**). To standardize comparisons, we defined a carbonyl *“rigidity index” R* that is independent of the number of CO residues that are labeled. For a given sample, index *R* is calculated as the ratio of experimental intensities of CO in CP-MAS and PARIS spectra normalized by the total signal intensity in the corresponding spectrum (equation S2).

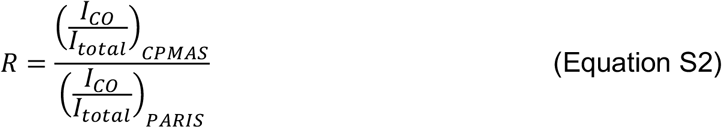

The *R* index gives values smaller than 1 if carbonyls are in environments of high mobility while higher values reflect an overall increase in rigidity. We obtained *R* values of 2.58, 0.84 and 5.00 for the recBVPrP^Sc^ (sample **A**), recBVPrP amyloid (sample **B**), and the HET-s(218-289) prion (sample **C**), respectively (Table S2). Although this *R* index is just a relative estimation and must be interpreted with caution, it indicates that overall, the three Phe residues in recBVPrP^Sc^ are not in environments of high mobility.

**Table S2.** Integrals (*I*) of the carbonyl (CO) peak of the 1D ^13^C CPMAS and PARIS spectra and carbonyl *Rigidity Index (R)*.

|  | **sample** | **( *I*<sub>CO</sub> / *I*<sub>total</sub> )<sub>CPMAS</sub>** | **( *I*<sub>CO</sub> / *I*<sub>total</sub> )<sub>PARIS</sub>** | ***R*** |
| --- | --- | --- | --- | --- |
| **A** | (<sup>13</sup>CO-Phe)-recBVPrP<sup>Sc</sup> | 0.270 | 0.105 | 2.58 |
| **B** | (<sup>13</sup>CO-Phe)-recBVPrP(23-231) amyloid | 0.170 | 0.201 | 0.84 |
| **C** | (<sup>13</sup>CO-Tyr)-HET-s(218-289) | 0.311 | 0.062 | 5.00 |

